# Elevated nonlinearity as indicator of transition to overexploitation in fish stocks

**DOI:** 10.1101/051532

**Authors:** Vasilis Dakos, Sarah M. Glaser, Chih-hao Hsieh, George Sugihara

## Abstract

Ecosystems may experience abrupt changes such as species extinctions, reorganisations of trophic structure, or transitions from stable population dynamics to strongly irregular fluctuations. Although most of these changes have important ecological and at times economic implications, they remain notoriously difficult to detect in advance. Here, we use a Ricker-type model to simulate the transition of a hypothetical stable fisheries population either to irregular boom-bust dynamics or to overexploitation. Our aim is to infer the risk of extinction in these two scenarios by comparing changes in variance, autocorrelation, and nonlinearity between unexploited and exploited populations. We find that changes in these statistical metrics reflect the risk of extinction but depend on the type of dynamical transition. Variance and nonlinearity increase similarly in magnitude along both transitions. In contrast, autocorrelation depends strongly on the presence of underlying oscillating dynamics. We also compare our theoretical expectations to indicators measured in long-term datasets of fish stocks from the California Cooperative Oceanic Fisheries Investigation in the Eastern Pacific and from the Northeast Shelf in the Western Atlantic. Our results suggest that elevated variance and nonlinearity could be potentially used to rank exploited fish populations according to their risk of extinction.

## Introduction

Ecosystem management is traditionally based on mechanistic models that describe and attempt to predict ecosystem dynamics. These models are hypotheses that represent our limited mechanistic knowledge and have been notoriously poor at prediction [1,2]. This shortcoming is evidenced by the fact that ever more elaborate models do not necessarily improve out-of-sample prediction of complex ecosystem dynamics [3,4]. This situation is especially evident in fisheries models where variables that are believed to be influential, such as temperature, can actually reduce predictability when included in the classic extended Ricker-model [1]. Even so, such models are still commonly used to set harvest guidelines [5]. There is an urgent need for improved warning methods for detecting imminent changes as ecosystems are exposed to novel stressors that create conditions for crossing thresholds at which unexpected and irreversible ecological changes might occur [6].

Fisheries present one of the most challenging cases for coping with high response uncertainty [7,8]. Fish populations are characterized by infamously unpredictable fluctuations driven by nonlinear dynamics [9], and magnified by environmental stochasticity [10]. This gives rise to stochastic chaos [11], a mathematical phenomenon that explains how environmental uncertainty can be amplified in a nonlinear fashion in the biological response [12]. Perhaps in a similar way, environmental changes or climate fluctuations have caused boom-and-bust cycles in sardines and anchovies [13] and have triggered regime shifts in a number of Pacific and Atlantic fish populations [14,15]. Boom-and-bust cycles and abrupt shifts to low population densities increase the risk of stochastic extinction, which has been documented on a global scale as a result of increasing fishing pressure [7]. The collapse of Atlantic cod in the early 1990s is a textbook example of such a shift due to overexploitation [16,17]. Size-selective fishing might have even left a signature on the life-histories of exploited populations [18]. Fishing-induced changes in demographic traits can theoretically trigger boom-and-bust dynamics in a population [9] (figure 1 a). Under such irregular fluctuations, the risk of stochastic extinction increases and forecasting this risk becomes more of a challenge.

**Figure 1.**
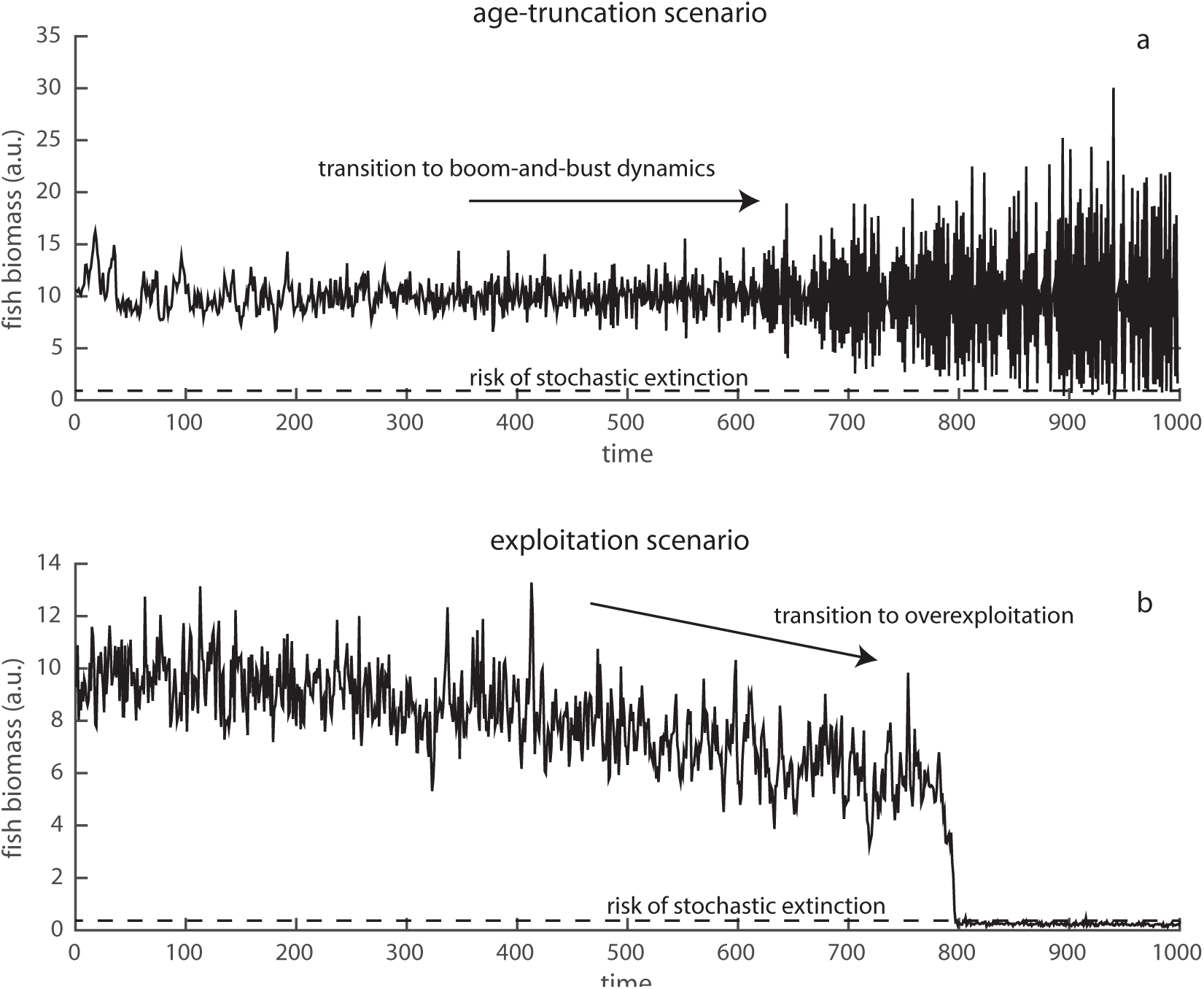
Two hypothetical trajectories of a fish stock under changing conditions. (a) Age-truncation effect: a gradual increase in population growth rate due to size-selective fishing leads to a shift from stable dynamics to boom-and-bust cycles (a combined effect of oscillations, chaos, and stochasticity that raises the risk of extinction during the bust phases). (b) Exploitation effect: a gradual increase in fishing can cause the population to shift rapidly to an overexploited state.

Clearly, coping with such a high level of response uncertainty requires alternative modeling strategies that go beyond traditional approaches [5,19]. One set of such strategies focuses on early warning signs that attempt to detect abrupt transitions based only on the statistical properties of the observed time series without requiring a specific mechanistic model [20]. Take the case of a stable fish population that, under increasing fishing pressure, might experience an abrupt shift in status from healthy to overexploited (figure 1 b). In this case, there are well understood changes in statistical properties of the time series that can be used to infer the risk of the approaching transition [20]. Such changes are caused by critical slowing down (CSD), a simple phenomenon by which stable systems close to local bifurcation points respond slowly to disturbances [21,22], and the statistical indicators that arise as corollaries are called critical slowing down (CSD) indicators. Bifurcation points are thresholds where the stability properties and, thus, the behavior of a system changes–typically through manipulation of a control parameter, such as harvesting or growth rate [23]. In particular, increasing variance and autocorrelation along a time series can signal a system’s proximity to an abrupt transition in ecosystem state. Thus, rising autocorrelation has been shown to be an indicator of the increasing risk of extinction in stable laboratory experiments with yeast [24] and zooplankton populations [25], whereas increasing variance marked the transition of lake dominance structure from piscivorous to planktivorous fish [26,27].

While the statistical early-warning indicators are model-free, they posit stable dynamics. Parallel work has focused on forecasting non-equilibrium and chaotic population dynamics that arise mechanistically from unstable attractors based on an approach called Empirical Dynamical Modeling (EDM) [1,28]. Similar to the statistical approach taken in CSD indicators, this approach is also equation-free. EDM is based on reconstructing an attractor manifold directly from time series ^1^. This reconstruction can be done with multiple time series of interest or with lags of a single time series [29]. In either case, how well the attractor is reconstructed can be verified by the ability to forecast future states. In fact, EDM has been shown to outperform equation-based approaches at forecasting recruitment in Sockeye salmon populations [1], dynamics of Pacific sardines [30], and the fate of experimental flour beetle populations [31]. With its capacity to forecast the future state of a system, EDM could be a useful approach to anticipate critical transitions.

Theoretically, when a dynamical system is approaching a critical transition (i.e. a bifurcation), stochastic events may more easily push a system across attractor boundaries, or far from equilibrium where dynamics may be also affected by a different attractor. This implies that, close to a bifurcation, the realized system dynamics may become increasingly state-dependent. State-dependence means that the future evolution of a system is determined by its current state. For example, approaching the boundary of the critical transition in the over-harvesting case (figure 1 b), dynamics will be increasingly affected by the alternative exploited and overexploited stable attractors. Therefore, close to the critical transition, forecasting the future system state requires knowledge of the current system state (i.e. local state information is critical). In contrast, far from the critical transition, there is only one stable state. In that case, local information is indistinguishable from global information. EDM can evaluate this state-dependence by comparing forecast performance obtained when using global versus local information to model the system [11,32]. If relying on local information gives a better forecast of system state compared to global information, the behavior of the system is deemed state-dependent. In principle, this concept can be generalized to more complex situations regardless of the type of attractors, such as bifurcations of cyclic attractors to chaos (e.g. figure 1a). In that way, while CSD indicators rely on changes in stability between stable states [33], EDM does not, so that in an unknown system, it is likely that the union of these two approaches may be more informative for anticipating critical transitions in ecosystems under stress.

Here, we study the behavior of EDM and CSD indicators along two different ecosystem transitions observed in time series that can increase the risk of stochastic extinction: the transition from stable equilibrium dynamics to irregular chaotic dynamics (figure 1 a), and the abrupt shift to an overexploited state (figure 1 b). Following former studies on nonlinear dynamics in fisheries [1,9,10,34], we generate time series using a stochastic Ricker type fishery model in which we assume a loss term due to fishing. We show how the behavior of nonlinearity, variance, and autocorrelation depends on the type of transition and discuss the capacity of these metrics as early warnings of loss in ecosystem stability. Finally, we measure these indicators on two long-term datasets of fishes from the Southern California Current ecosystem and U.S. Northeast Shelf system in the North Western Atlantic. Our aim is to illustrate how these model-free approaches can be helpful for ecosystem management by quantifying resilience across populations.

## Materials and Methods

### Simulated data

We used a discrete Ricker type model that describes the logistic growth of population N to which we added a loss term due to fishing. The model reads:

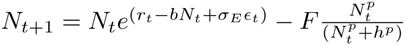
where *r*_*t*_ is the intrinsic growth rate, *b* defines the strength of density-dependence (= *r*_*t*_/*K*, where *K* is the carrying capacity set by the environment (=10)), and fishing follows a sigmoid functional response (p=2), with half-saturation *h* (=0.75) and maximum fishing rate *F*. We assumed process error in the model to represent environmental stochasticity with a Gaussian term *∊*_*t*_ of zero mean and σ_*E*_ (=0.25) standard deviation. We also considered demographic stochasticity in the growth rate *r*_*t*_ by using exponential filtering at each time step (*r*_*t*_ = *r*_0_*e*^(σ_*r*_*∊*_*r*,*t*_)^), where *r*_0_ is the mean and *σ*_*r*_ (=0.1) the standard deviation of the Gaussian noise term *∊*_*t*_ [35]. Although we used fisheries as an example, our analysis and results are generic to any ecosystem or population model that exhibits similar dynamics and transitions.

We considered two scenarios where fishing can potentially trigger dynamical changes in the behavior of a population. In the first scenario, we hypothesized that fishing leads to changes in population demographic traits due to age-truncation caused by size-selective removal (figure 1 a). Selective fishing of large individuals in a population may cause small size individuals to mature at an earlier age [18,36]. Earlier age-at-maturation can be associated with an overall increase in the intrinsic growth rates of exploited fish populations [9]. Here, we mimic this “age-truncation effect” by gradually increasing growth rate *r*_0_ (=[0.01, 3]), while setting the overall fishing rate *F* to zero. Increasing growth rate in the Ricker model exhibits a well-known series of transitions from stable equilibrium dynamics to period-doubling bifurcations that give rise to cycles and eventually chaos [37].

In the second scenario, we hypothesized that a fish population runs the risk of overexploitation due to direct fishing pressure (figure 1 b) [38]. We labeled this the “exploitation effect” scenario that we simulated by progressively increasing fishing rate *F* (r=[0, 3]), starting from a stable non-fluctuating population (*r*_0_ = 0.75). Intensifying fishing rate leads to the abrupt collapse of the population due to the crossing of a fold bifurcation that forces the population to shift from an underexploited to an overexploited state of low abundance.

For both age-truncation and exploitation scenarios, we gradually increased the bifurcation parameters, r0 and F respectively, in 100 equidistant steps. At each step, we burned-in the models for a period of 100 time steps to discard transients, and we simulated another 100 points to use for analysis. For each step of the bifurcation parameter, we produced 1000 replicate time series that were used to estimate nonlinearity and critical-slowing-down indicators. We also tested the behavior of the indicators while changing conditions. To do this, we continuously increased growth rate *r*_0_ (=[0.01, 3]) and fishing rate *F* (=[0, 3]) in 200 time steps using a sliding window of half size the time series (that is, 100 points). We reported indicator means and 95% confidence intervals based on the 1000 replicates.

### Empirical fish data

We used fish data collected from scientific surveys on the Northeast Shelf (NES) in the northwest Atlantic and in the southern California Current Ecosystem (CCE) in the eastern Pacific. NES data were collected through the Northeast Fisheries Science Center. These data are relative biomass estimates generated from an annual fall bottom trawl survey and include 29 stocks of demersal fishes sampled from 1963 to 2008. Of these, 20 stocks were exploited (subject to fishing pressure). CCE data were collected through the California Cooperative Oceanic Fisheries Investigations (CalCOFI). These data are relative biomass estimates generated from regular ichthyoplankton tows and include 29 coastal-neritic fish species [39] sampled from 1951 to 2007. Among the 29 species, 16 were exploited and 13 were unexploited. The NES data are available from the Northeast Fisheries Science Center, National Marine Fisheries Service (NMFS), National Oceanic and Atmospheric Administration (NOAA), USA (http://www.nefsc.noaa.gov/nefsc/saw/). The CCE fish data are available from the Southwest Fisheries Science Center, NMFS, NOAA, USA (http://coastwatch.pfeg.noaa.gov/erddap/search/index.html?page=1&itemsPerPage=1000Searc

### Critical Slowing Down indicators

Critical slowing down (CSD) is defined as the decrease in recovery rate upon small perturbations in the vicinity of local bifurcation points [23]. It is a generic property of dynamical systems that undergo transitions between different attractors when a stress parameter crosses a threshold. In mathematical terms, CSD is associated with a diminishing dominant eigenvalue λ, where λ defines the rate of exponential decay of a perturbation close to equilibrium (Δ*x* = *e*^−λ*t*^) [21]. The consequence of this slow decay is that both variance and autocorrelation of the recorded ecosystem dynamics will increase close to a transition point [20]. We estimated variance as coefficient of variation (*CV* = standard deviation/mean), and autocorrelation at lag-1 (*AR*1) as the Pearson correlation for lagged time series at one step [40].

### Nonlinearity indicators derived from Empirical Dynamical Modeling

We propose to use nonlinearity (i.e. quantification of state-dependence) as an additional indicator for anticipating critical transitions. To determine whether a time series reflects linear or nonlinear processes, we compared the out-of-sample forecast skill of a linear model (i.e. relying on global information) versus an equivalent nonlinear model (i.e. relying on local information). This involves state space reconstruction (aka EDM) using lagged coordinate embeddings with a two-step procedure as follows. First, we used simplex projection [28] to determine the embedding dimension (*E*) of the system where E represents the number of independent variables needed to reconstruct the system state-space and is operationalized as the number of lagged coordinates used to reconstruct the system attractor. Second, using this embedding dimension, we used S-map [11] to compare linear versus nonlinear forecasting models, by tuning a nonlinear weighting parameter *θ*. If the forecast skill of the nonlinear model (*θ* > 0) outperforms that of the linear model (*θ* = 0), the observed dynamics in the system are classified as nonlinear (or state dependent). Forecast skill is evaluated based on the correlation between S-map predicted out-of-sample values and the actual observed values in the time series. See a suite of articles fully describing this established methodology [1,32,41,42].

We applied the above procedure after first-differencing and standardizing both simulated and fisheries time series [32]. As EDM requires a time series of at least 30 observations [43], we used simulated time series of 100 time steps and selected empirical fish records that contained at least 30 points. We produced simplex projections using lagged coordinates of one time step (*τ* = 1) for a series of different embedding dimensions *E* (1 through 10 for the fish data and 1 through 3 for simulated data - we used a smaller range of *E* for the simulated data as we knew a priori the dimensionality of the Ricker model attractor following Whitney’s theorem that *n* ≤ *E* ≤ 2*n* + 1 where *n* is the dimensionality in the systems (*n* = 1 for our model)). We applied a cross-validation approach, using *E* + 1 vectors, to estimate Pearson correlation (*ρ*) between observed and forecast values to choose the best embedding dimension *E* for the system. The best *E* was then used for fixing the embedding space in the S-map. In the S-map we did the same cross-validation as above but using all vectors (not just nearest neighbors) to perform the forecasting. The vectors, however, are weighted exponentially according to the tuning parameter *θ*. When *θ* = 0, all vectors are weighted equally (linear case). When *θ* > 0 (nonlinear case), neighbors closer to the predicted vector become more important for forecasting (the increasingly smaller neighborhood thus represents increasing state-dependence). We tried a range of *θ* values [0, 0.001, 0.005, 0.01, 0.05, 0.1, 0.2, 0.3, 0.5, 1, 2, 5], and for each *θ* value we compared the improvement in forecasting skill.

We estimated the improvement in forecasting skill of the nonlinear over the linear model as the difference in Pearson correlation: Δ*ρ* = *max*(*ρ*_*θ*_—*ρ*_*θ*=0_): the maximum difference between the correlation *ρ*_*θ*_ at each *θ* to the correlation *ρ*_*θ*_ = 0 found for *θ* = 0. In other words, Δ*ρ* defines how much the nonlinear model (*θ* ≠ 0) outperformed the linear model, which is used to quantify nonlinearity [9]. We therefore use the metric of Δ*ρ* as indicator of nonlinearity.

All simulated data and the estimation of the nonlinearity and CSD indicators were produced using MATLAB v2015a (Mathworks). For open source options, nonlinearity indices can be estimated using the rEDM package (https://github.com/ha0ye/rEDM), and CSD indicators can be computed using the R package earlywarnings (https://github.com/earlywarningtoolbox). For the fish time series, missing values were omitted in estimating the indicators.

## Results

In the age-truncation scenario (figure 2 a), a gradual increase in growth rate in the Ricker model caused a sequence of transitions from stable to excitable dynamics that led to regular oscillations and then chaos (figure S1a). The boundaries between different attractors were difficult to define in the presence of stochasticity. Nonetheless, we clearly observed different behaviors in *CV*, *AR*1, and Δ*ρ* that were distinct in each dynamic regime defined by the deterministic model (figure S1b). Before the onset of spiraling stable dynamics, variance, autocorrelation, and nonlinearity all decreased (figure 2 b). Interestingly, Δ*ρ* decreased slightly even after the onset of the spiraling stable dynamics. Close to the onset of cycles, *CV* and Δ*ρ* increased but AR1 kept decreasing. Approaching the transition to chaos and inside the chaotic regime, *CV* and Δ*ρ* continued to increase, and so did *AR*1 but much more weakly (figure 2 b). Overall, only the relationship between *CV* and Δ*ρ* was consistent across all regimes. *AR*1 was sensitive to the presence of oscillations, as the spiraling and cyclic attractors shifted correlations from positive to negative values (figure S2b).

**Figure 2.**
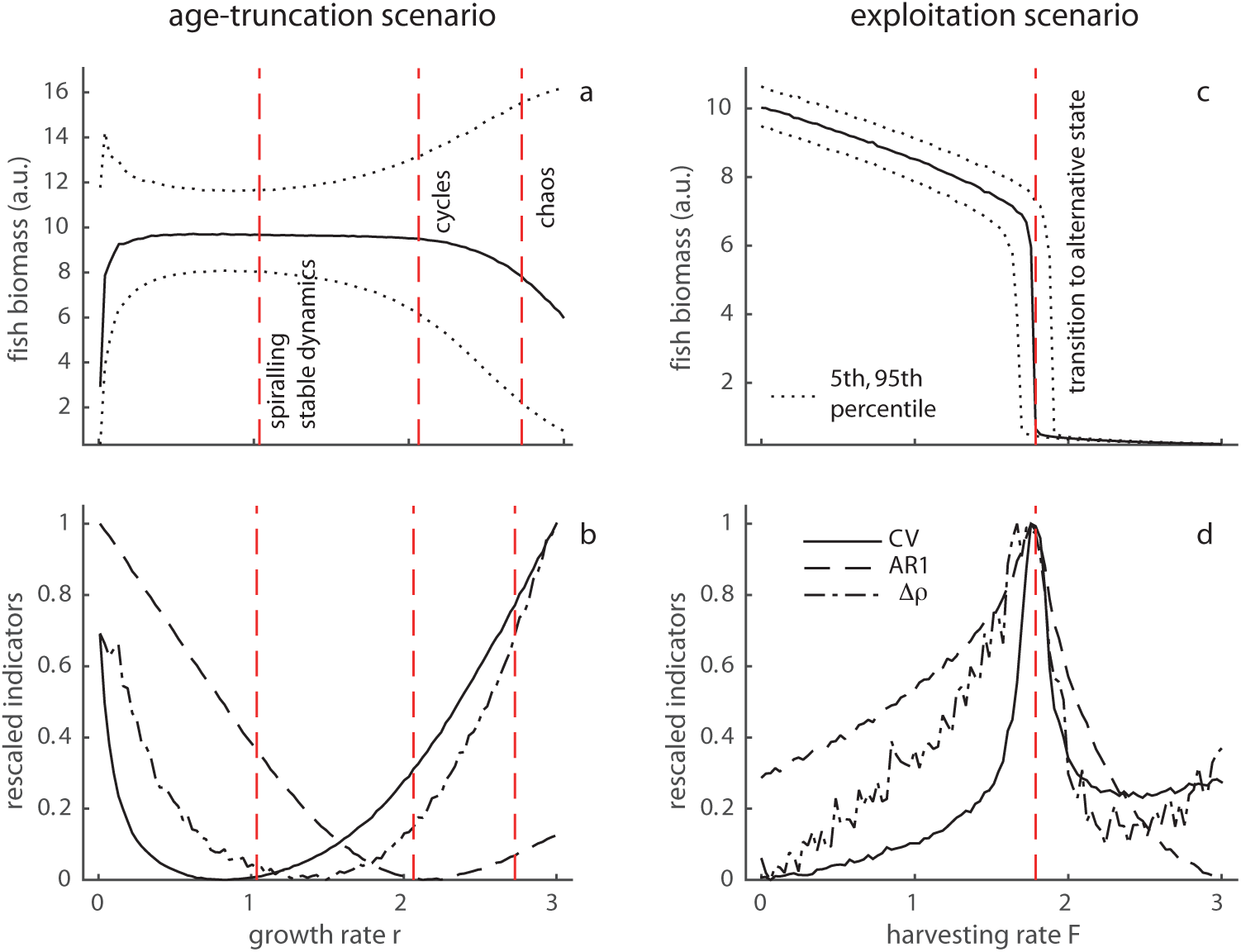
Age-truncation scenario: (a) median value (of 1000 replicates) of mean (over time, per replicate series) fish abundance as a function of growth rate *r*. Red dashed lines indicate the thresholds between dynamics that are stable, spiraling stable, cycles, and chaotic of the deterministic model. (b) Rescaled coefficient of variation *CV*, autocorrelation at-lag-1 *AR*1, and nonlinearity index Δ*ρ* (means of 1000 replicate time series) as a function of r. Exploitation scenario: (c) median value (of 1000 replicates) of mean (over time, per replicate series) fish abundance as a function of fishing rate F. Vertical dashed line indicates the threshold at which 50% of the 1000 populations collapse to the alternative overexploited state. (d) Rescaled *CV, AR*1, and Δ*ρ* (means of 1000 replicate time series) as a function of *F*.

In the exploitation scenario, a gradual increase in fishing rate *F* caused our model fish populations (mean biomass thereof) to experience a slight decrease until populations suddenly collapsed to an overexploited state (figure 2 c). Due to the process error in the model, overexploitation occurred earlier than the actual fold bifurcation of the deterministic model (figure S1b). Nonetheless, both the CSD indicators (variance *CV*, autocorrelation *AR*1) and the EDM-based indicator (nonlinearity Δ*ρ*) strongly increased before the collapse (figure 2 d). Once the transition was crossed, the pattern reversed: *CV, AR*1, and Δ*ρ* dropped, marking the progressive gain in resilience and decrease in nonlinearity of the overexploited state.

Ideally, for populations that have a history of monitoring, a manager would have available population abundance estimates or growth rates at different levels of fishing pressure in order to estimate these three indicators. In reality, however, fishing and any other environmental conditions all change continuously. Therefore, we studied this more realistic scenario by measuring CSD and nonlinearity indicators along a time series at continuously changing conditions (figure S3) using a sliding window approach [40]. We found similar patterns to those obtained in the case of stationary distributions (figure 2), but with strong uncertainties especially in the case of nonlinearity (figure S3g, h).

We tested our theoretical predictions on empirical fish time series. In our fisheries records, however, we cannot vary F as done in the simulations, but we can only discriminate populations based on whether they were commercially fished (i.e. exploited) or not (i.e. unexploited or bycatch). Using this discrimination as a proxy for fishing pressure, we tested for differences in *CV, AR*1, and Δ*ρ* between exploited versus unexploited populations (figure 3). We found that mean Δ*ρ* and *AR*1 were higher for exploited populations in both datasets (figure 3 b, c), whereas mean *CV* was higher in exploited populations in the CCE but not in the NES dataset (figure 3 a). However, only Δ*ρ* and *CV* were significantly higher in the CCE exploited populations (t-test with unequal variances). Taking both datasets together (table S1), we found negative correlations between *CV* and *AR*1 for both exploited and unexploited populations, stronger positive correlation between *CV* and Δ*ρ* for unexploited than exploited populations, and also stronger positive correlation between *AR* and Δ*ρ* for unexploited than exploited populations. However, similar to our simulated time series, only the correlations between *CV* and Δ*ρ* were significant.

**Figure 3.**
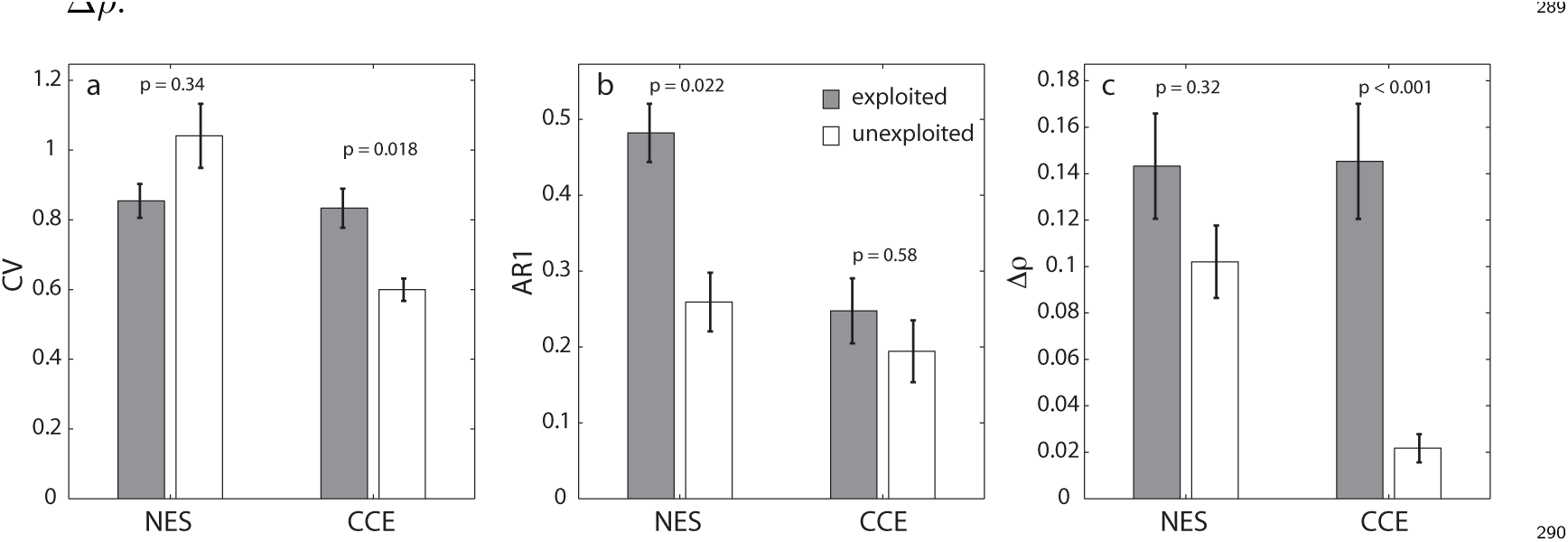
Critical slowing down and nonlinearity indicators in empirical time series from the North Eastern Shelf (NES) and California Current Ecosystem (CCE) fish stocks. (a) Coefficient of variation *CV*, (b) autocorrelation at-lag-1 *AR*1, (c) nonlinearity index Δ*ρ*. Grey bars represent indicators for exploited populations (targeted by commercial fisheries); white bars represent indicators for unexploited populations (not fished or bycatch). Bars reflect mean values with standard error 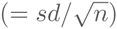, and *p* values are derived from *t*-tests with unequal variances between populations.

The patterns between *CV, AR*1, and Δ*ρ* that we observed in our empirical fisheries data generally matched the theoretical patterns we derived from our simulated time series (figure 4). We approximated simulated populations as exploited or unexploited by grouping time series far from and close to the collapse for the exploitation effect scenario (unexploited: 0 < *F* ≤ 0.97, exploited: 0.97 <F< 1.79), and before and after the deterministic limit of cyclic dynamics in the age-truncation effect scenario (unexploited: 0.01 < *r*_0_ ≤ 2.06, exploited: 2.06 < *r*_0_ < 3). In both scenarios *CV* and Δ*ρ* were higher in the exploited than in the unexploited populations, while *AR*1 was higher for the exploited populations in the exploitation scenario (figure 4 b), but lower in the age truncation scenario (figure 4 a). Overall, the correlations between *CV* and Δ*ρ*, and between *CV* and *AR*1, matched the empirical relationships, but not between *AR*1 and Δ*ρ*.

**Figure 4.**
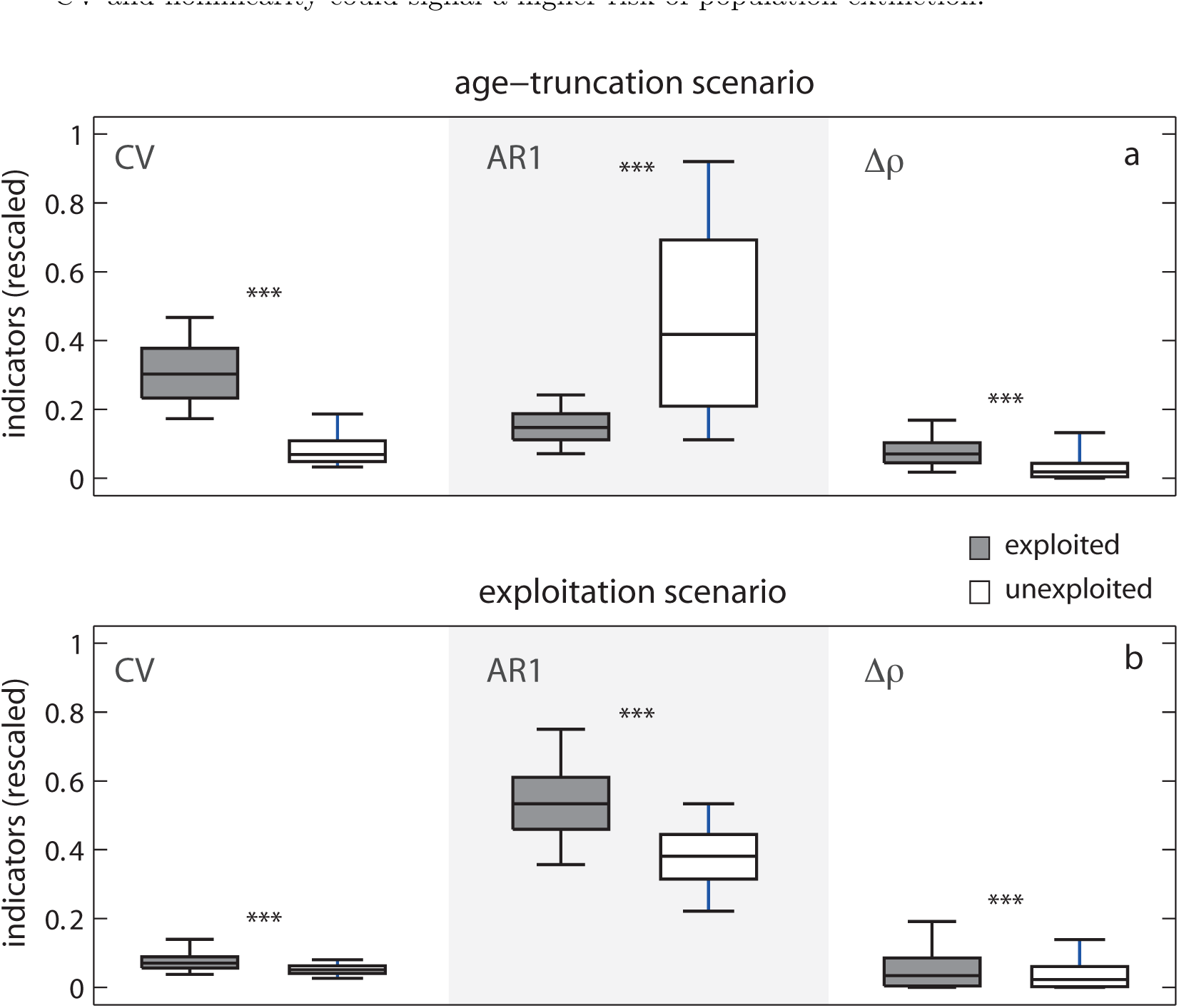
Distributions of critical slowing down and nonlinearity indicators (boxplots median, 5 and 95 percentiles). (a) Age-truncation scenario. We categorized populations as unexploited or exploited based on their growth rates: 0.01 < *r* ≤ 2.06 (unexploited), 2.06 < *r* < 3 (exploited). We assumed that populations were exploited if their growth rates exceeded the limit (*r* ≃ 2.06) that corresponds to the onset of cyclic dynamics in the deterministic model, as we assumed that age-truncation due to fishing leads to irregular oscillations. (b) Exploitation scenario. We categorized unexploited and exploited populations before the collapse to overexploitation (*F* ≃ 1.79) based on fishing rate: 0 < *F* ≤ 0.97 (unexploited), 0.97 < *F* < 1.79 (exploited). Grey bars are exploited populations; white bars represent unexploited populations. *** indicates *p* < 0.001 from t-tests with unequal variances between populations.

## Discussion

The nonlinear dynamics of most ecological systems have puzzled ecologists for a long time. Driven by chaotic dynamics, stochasticity, or alternative attractors, our ability to accurately describe nonlinear systems and to efficiently manage them remains limited. Here, we explored whether Empirical Dynamic Modelling and Critical Slowing Down indicators can be used for detecting changes in ecological dynamics using fisheries as an example. These two approaches belong to a class of model-free methods for describing ecological dynamics. EDM has never been explored as a means of measuring ecosystem stability or detecting the risk of dynamical transitions in the generic sense as CSD indicators do. Our results show that elevated nonlinearity Δ*ρ* based on EDM can be a novel indicator for the proximity to dynamical transitions between alternative states as well as to transitions from stable dynamics to irregular oscillations.

Bifurcation analysis reveals that the indicator patterns we observed capture the underlying deterministic stability of the Ricker model (figure S1). In the age-truncation scenario, changes in growth rates move the system from a single stable state to multiple cyclic states and finally to chaos (figure S1a). In the exploitation scenario, increasing fishing pressure brings the system close to the fold bifurcation where the stable attractor merged with the unstable saddle and propels the system to the alternative state (figure S1b). In both scenarios environmental and demographic stochasticity forces the system across these dynamical regimes. As a result, trajectories are increasingly affected by the stability properties of the different attractors. EDM captures this increasing state-dependence in the system dynamics, as it can better follow future trajectories when it considers the local information of state space. Increasing variance also reflects the generic rising divergence in the state space due to the existence of multiple states. Autocorrelation at lag-1, however, is sensitive to the type of dynamics between the two scenarios: it consistently rises in the exploitation scenario, but builds-up or breaks down under age-truncation. In that case, it might have been more informative looking at higher spectra than first lags.

Elevated nonlinearity and variability in stressed populations are the most consistent patterns in both scenarios we tested. This observation actually implies that these indicators may be better suited to detect changes in the dynamics observed in natural populations. It also implies that the source of these changes goes beyond the loss of stability across bifurcation points. More generally, rising variance and nonlinearity can be understood as a statistical phenomenon of increasing state-dependence [11] that does not require the existence of stable attractors. In reality, population dynamics are typically the result of a mix of transients across stable and unstable attractors affected by environmental and demographic stochasticity. CSD indicators can capture changes in these dynamics but only when it comes to stable equilibria in the presence of weak stochasticity [20]. Identifying transitions across chaotic attractors or, more generally, in systems with nonlinear dynamics may be difficult with CSD indicators [44,45]. Thus, EDM-derived nonlinearity is broader in its application, and it can capture changes in dynamics beyond stable attractors typical of the dynamics encountered in natural systems.

Based on our findings, a resource manager could estimate both CSD and EDM metrics in order to infer levels of stress and rank populations according to their potential risk to extinction [46]. For instance, Krkosek and Drake [47] looked at patterns of *CV* and *AR*1 for Pacific salmon populations and found that they were higher for pink salmon stocks that had a population growth rate close to zero. In this work, the authors assumed that salmon populations would suffer a transcritical bifurcation due to growth rates approaching zero. Trends in *CV* and *AR*1 in our analyses are in line with these findings (figure 2). Moreover, we also find that nonlinearity increases at decreasing growth rates. Our results imply that regardless of the type of transition (be it a transcritical, a fold, or a period-doubling bifurcation), a simultaneous increase in CV and nonlinearity could signal a higher risk of population extinction.

Trends in empirical data generally agreed with our simulated results. We found elevated nonlinearity in exploited fish populations for both NES and CES datasets, higher variability only for the CCE dataset, and a stronger AR1 in exploited populations for both datasets (figure 3). These results imply that some populations were under the exploitation stress scenario, while others followed the age-truncation scenario (figure 4). It is hard, though, to identify which scenario mostly affected each set of populations (table S1). The only significant result is that, when taking all populations together, *CV* and nonlinearity show a consistent positive relationship (table S1).

While we compared populations sampled at the same geographical ranges and under similar climatic conditions, it is still difficult to draw strong conclusions from these patterns. First, we assumed that exploited populations are affected by both overexploitation and age-truncation. Second, we did not take into account life-history traits or stochastic events that may also affect population dynamics. Pinsky et al. [7] demonstrated that in addition to fishing, environmental stochasticity and high growth rates can affect the risk of collapse in fish populations. In our analysis, we used age-at-maturation as proxy for population growth rate and we assumed that age-truncation pushes populations to reproduce faster [9]. We found negative (but non-significant) correlations between nonlinearity and variance versus age-at-maturation (figure S4), which hints that faster growth rates may lead to stronger irregular fluctuations and a higher risk of collapse.

Of course, critical slowing down and nonlinearity indicators are not bulletproof metrics. We found strong fluctuations in nonlinearity estimates, especially in the exploitation scenario (figure S2c, d). Changes in variance and autocorrelation can also be unreliable in the presence of short time series [40], high levels of stochasticity [48], fast changing stress drivers [45,49], or due to portfolio effects [47] and life-history strategies [50]. Further research is needed to find if similar constraints hold for the elevated nonlinearity indicators we proposed here.

In the quest for understanding and anticipating future ecosystem responses, testing novel and alternative approaches is of high priority. The indicators we examined here contribute to equation-free, data-driven approaches that aim at quantifying differences in the resilience of populations under increasing environmental stress. Translating such differences to a risk assessment scheme might be a useful tool for improving ecosystem management in the face of global environmental change.

## Acknowledgments

We would like to thank Florian Grziwotz and Arndt Telschow for valuable comments on the manuscript. We also thank the California Cooperative Oceanic Fisheries Investigations, Southwest Fisheries Science Center, and Northeast Fisheries Science Center for the use of the data. Hui Liu assembled the NES data used in the study. VD acknowledges support by a MarieCurie EU IntraEuropean fellowship (2013-2015) and funding from the Center for Adaptation to a Changing Environment (ACE) at ETH Zürich. CHH is supported by the National Center for Theoretical Sciences, Foundation for the Advancement of Outstanding Scholarship, and Ministry of Science and Technology of Taiwan.

---

**Table S1.**
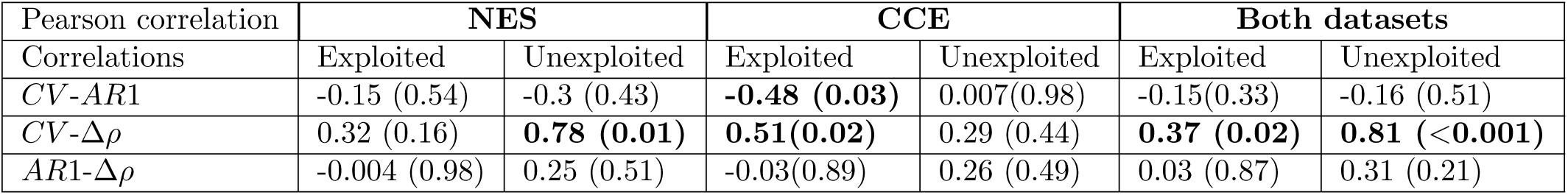
Pearson correlations between indicators for exploited and unexploited populations in the Northeast Shelf System (NES) and the southern California Current Ecosystem (CCE). *p*-values are given in parentheses and those < 0.05 are bolded

**Figure S1.**
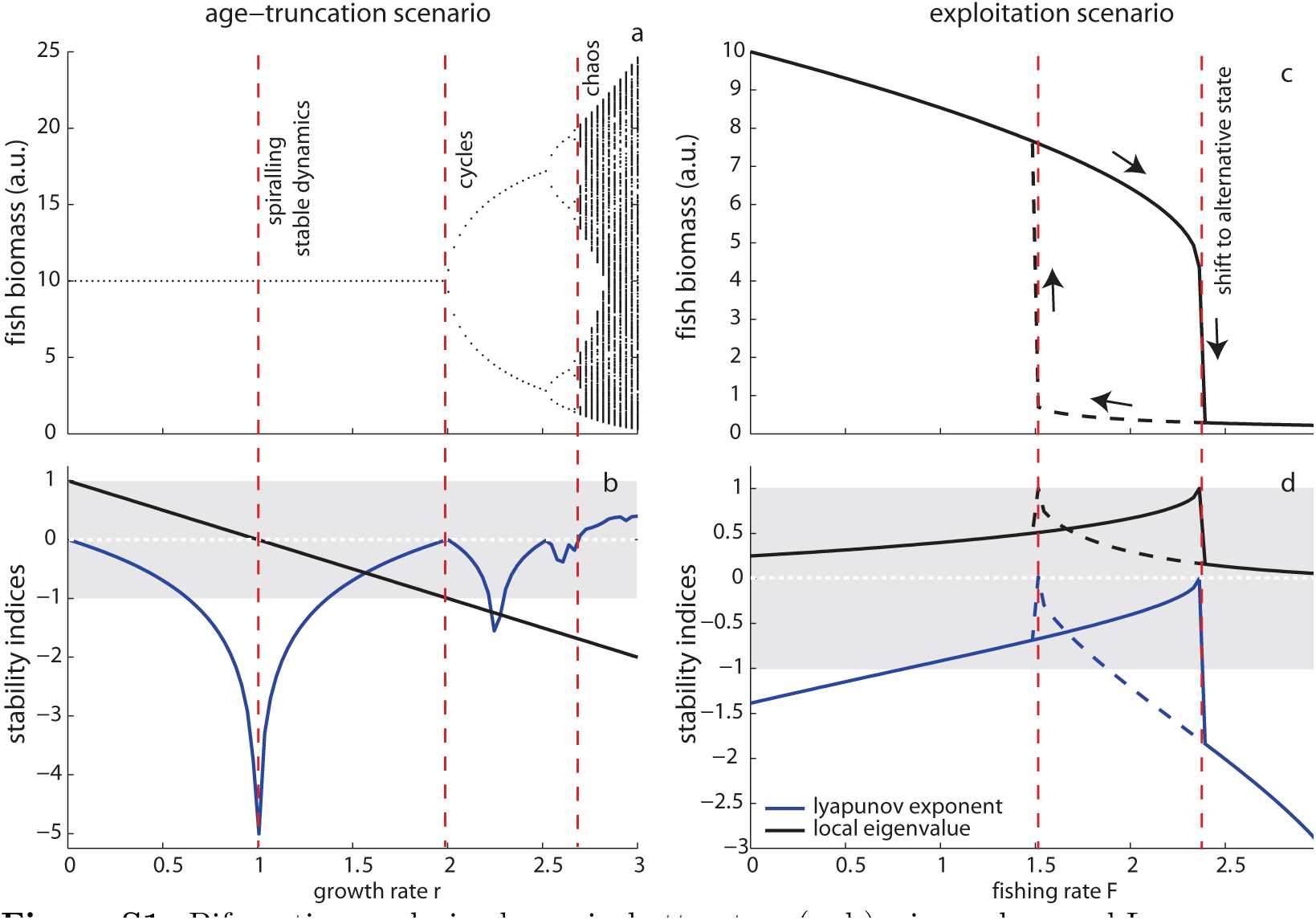
Bifurcation analysis, dynamical attractors (a, b), eigenvalues and Lyapunov exponents (c, d) of the deterministic model. In (a, b) we assume that age-truncation shifts the basic reproductive rate to higher values (increase in growth rate r). Dynamically, a stable equilibrium is replaced by stable but spiraling dynamics, and as growth rate increases, the system starts to oscillate in cycles of increasing periods before becoming chaotic (b). In (c, d) we assume that fishing can increase harvesting pressure. Dynamically, we have a stable equilibrium that is replaced by an alternative state at the crossing of a fold bifurcation. Theoretically, we measure stability based on the eigenvalue (the rate of return to equilibrium) and the Lyapunov exponent (the rate of divergence from equilibrium after a small perturbation). Eigenvalues crossing unity (−1, +1) signify loss of stability, negative Lyapunov exponents signify convergence (stability), and positive Lyapunov exponents signify divergence (instability and chaotic dynamics).

**Figure S2.**
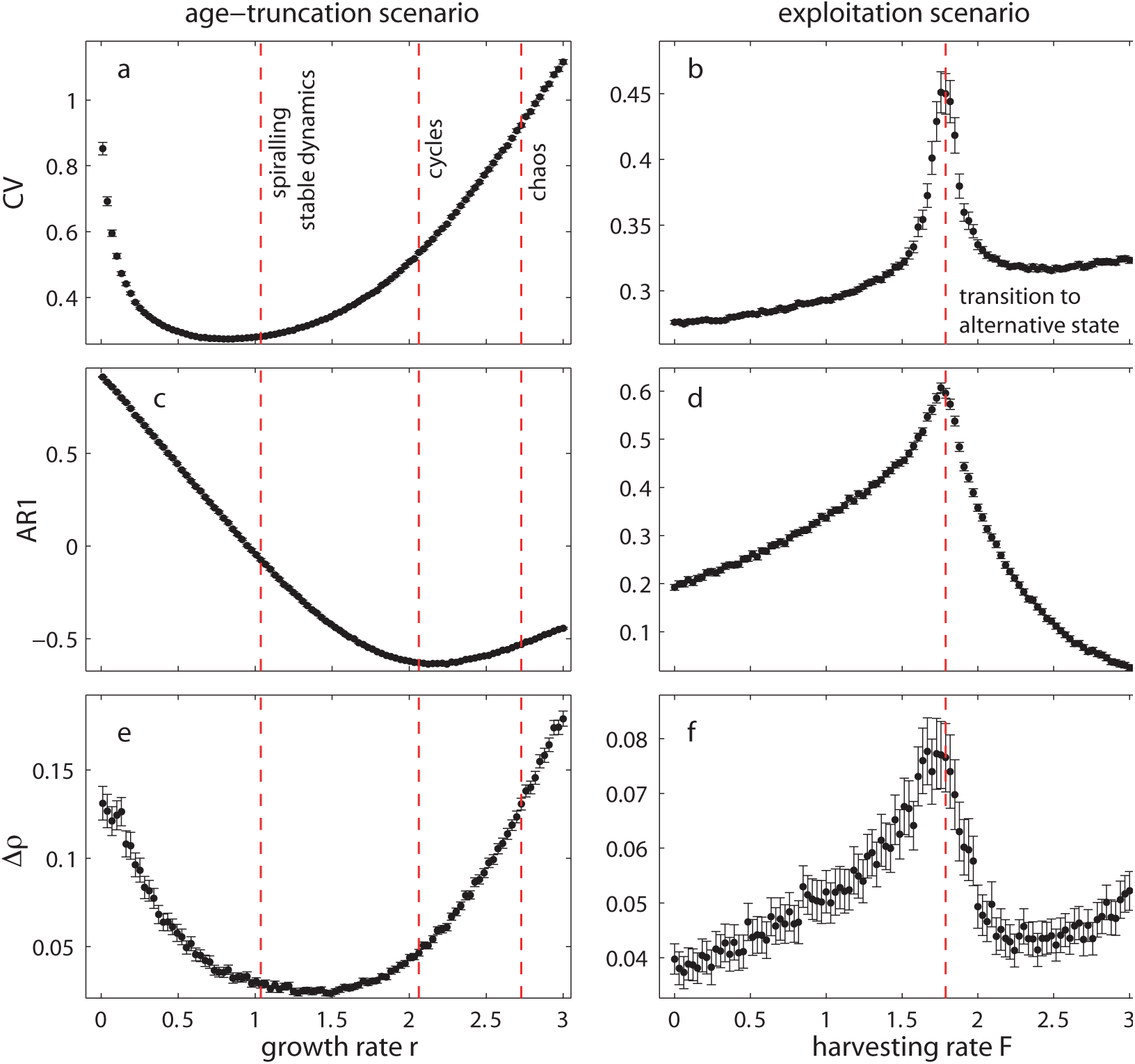
Critical slowing down and nonlinearity indicators for the two scenarios (mean and 95% confidence intervals (CI) based on 1000 simulations) for each level of fishing rate F and growth rate *r*. (a, c, e) In the age-truncation scenario, changes are not monotonic but depend on the type of dynamical regime. Red dashed lines indicate the thresholds between stable, spiraling stable dynamics, cycles, and chaos of the deterministic model. (b, d, f) All indicators change monotonically before the shift to overexploitation. Red dashed line indicates the threshold at which 50% of the 1000 populations collapse to the alternative overexploited state. Note the strong CI for Δ*ρ* in the exploitation scenario. Critical slowing down indicators: coefficient of variation *CV* and autocorrelation at-lag-1 *AR*1. Nonlinearity indicator Δ*ρ*.

**Figure S3.**
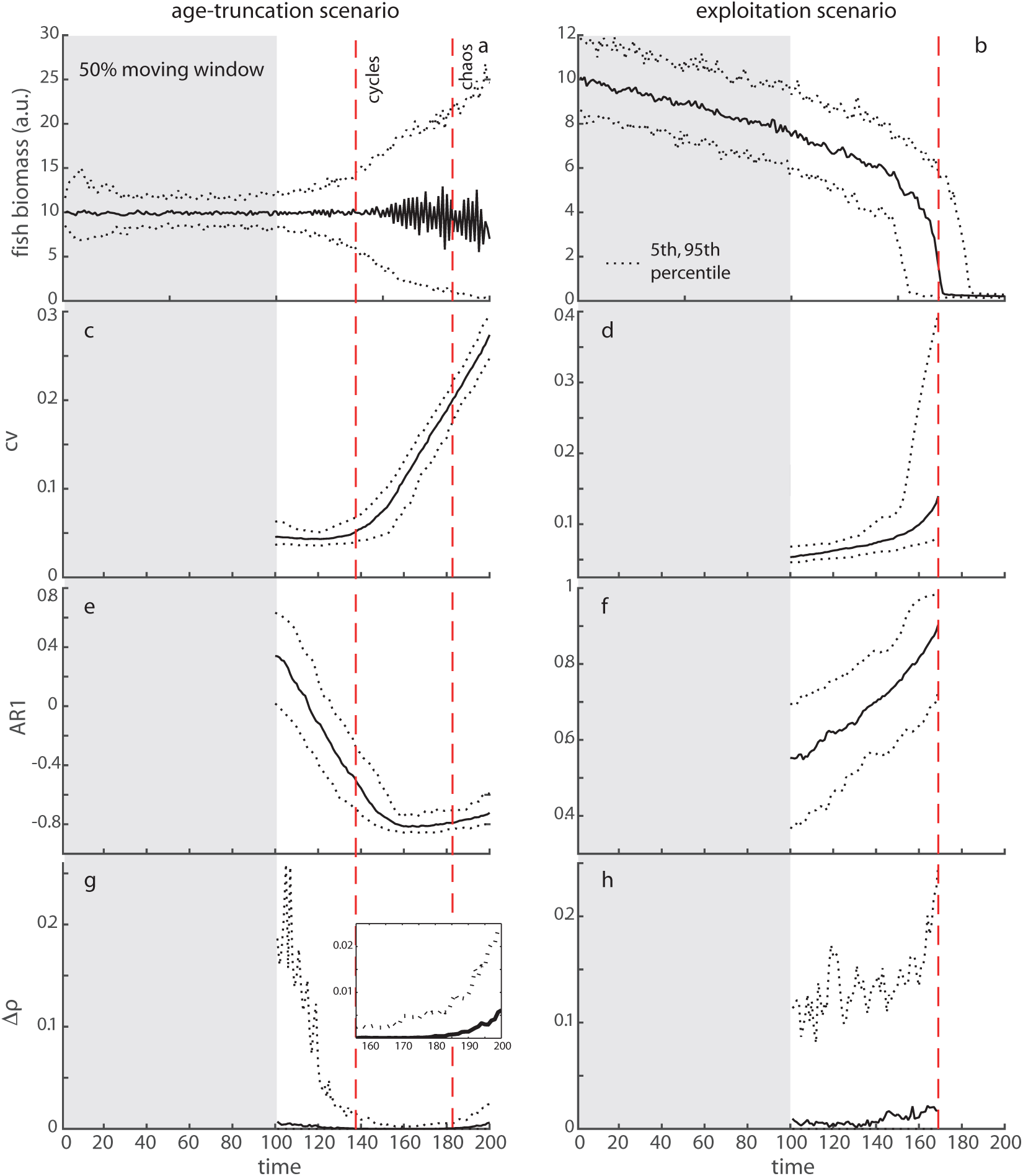
Critical slowing down and nonlinearity indicators estimated within a moving window of half the size of the time series. Time series are generated by continuously increasing growth rate r and fishing rate F respectively in 200 time steps. (a, b) Fish abundance (median, 5th and 95th percentiles based on 1000 simulations). (c, e, g) In the age-truncation scenario, changes are not monotonic but depend on the type of dynamical regime. Red dashed lines indicate the thresholds between cycles and chaos of the deterministic model. (d, f, h) In the exploitation scenario, all indicators change monotonically before the shift to overexploitation. Red dashed line indicates the threshold at which 50% of the 1000 populations collapse to the alternative overexploited state. Critical-slowing-down indicators: coefficient of variation *CV* and autocorrelation at-lag-1 *AR*1. Nonlinearity indicator Δ*ρ*.

**Figure S4.**
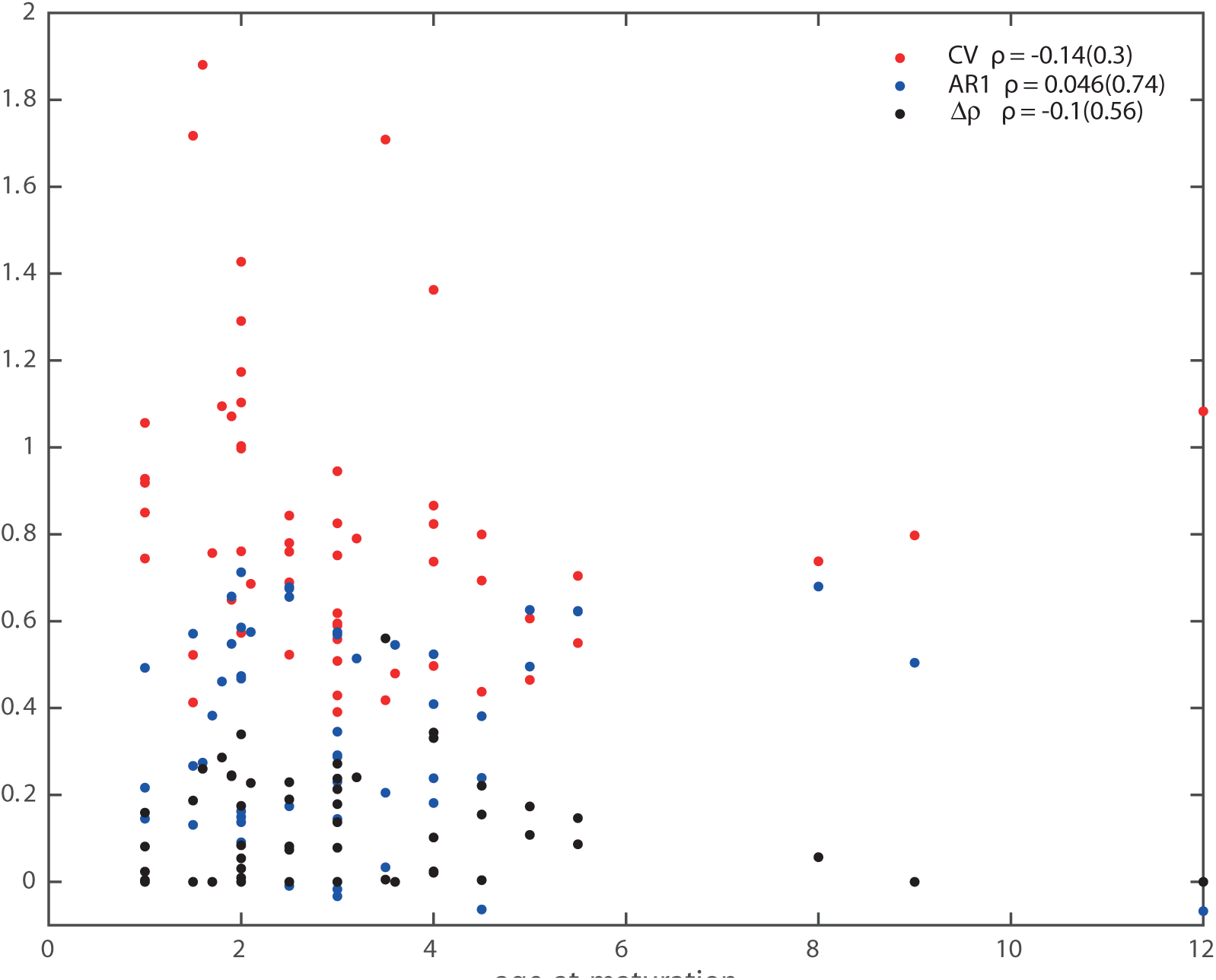
Relationship between critical slowing down and nonlinearity indicators versus age-at-maturation of southern California Current Ecosystem fisheries data. Age-at-maturation is considered as the proxy for growth rate. We found negative (albeit non-significant) correlations in *CV* and Δ*ρ*, but not *AR*1, versus age-at-maturation. This implies that faster growing populations are characterized by elevated nonlinearity and variability. (Pearson *ρ* correlation with significance p-value in parenthesis).

## Footnotes

1 https://www.youtube.com/watch?v=rs3gYeZeJcw

